# Integrative classification of human coding and non-coding genes based on RNA metabolism profiles

**DOI:** 10.1101/073643

**Authors:** Neelanjan Mukherjee, Lorenzo Calviello, Antje Hirsekorn, Stefano de Pretis, Mattia Pelizzola, Uwe Ohler

**Affiliations:** Berlin Institute for Medical Systems Biology, Max Delbrück Center for Molecular Medicine, Berlin, Germany; Center for Genomic Science of IIT@SEMM, Fondazione Instituto Italiano di Tecnologia, Milan, Italy.; Department of Biology, Humboldt University, Berlin, Germany; Department of Computer Science, Humboldt University, Berlin, Germany

## Abstract

The pervasive transcription of the human genome results in a heterogeneous mix of coding and long non-coding RNAs (lncRNAs). Only a small fraction of lncRNAs possess demonstrated regulatory functions, making it difficult to distinguish functional lncRNAs from non-functional transcriptional byproducts. This has resulted in numerous competing classifications of human lncRNA that are complicated by a steady increase in the number of annotated lncRNAs.

To address these challenges, we quantitatively examined transcription, splicing, degradation, localization and translation for coding and non-coding human genes. Annotated lncRNAs had lower synthesis and higher degradation rates than mRNAs, and we discovered mechanistic differences explaining the slower splicing of lncRNAs. We grouped genes into classes with similar RNA metabolism profiles. These classes contained both mRNAs and lncRNAs to varying degrees; they exhibited distinct relationships between steps of RNA metabolism, evolutionary patterns, and sensitivity to cellular RNA regulatory pathways. Our classification provides a behaviorally-coherent alternative to genomic context-driven annotations of lncRNAs.

**Highlights:**

- High-resolution 4SU pulse labeling of RNA allows for quantifying synthesis, processing and decay rates across thousands of coding and non-coding transcripts.
- Synthesis and processing rates of lncRNAs are lower than mRNAs, while degradation rates were substantially higher
- Differences in the splicing efficiency between slow/lncRNA and fast/mRNA introns are explained by GC-content, splicing regulatory elements and unphosphorylated RNA poll II.
- A new annotation-agnostic classification of RNAs reveals seven clusters of lncRNAs and mRNAs with unique metabolism patterns that provides behaviorally coherent subsets of lncRNAs.
- Classes are distinguished by evolutionary patterns and sensitivity to cellular RNA regulatory pathways.

## Introduction

Over the last decade, the increasing evidence for pervasive transcription of the human genome has spurred efforts to identify and functionally characterize long non-coding RNAs (lncRNAs) (Birney et al., 2007). lncRNAs are defined as non-coding transcripts longer than 200 nucleotides, an ad-hoc convention to differentiate them from well-characterized small non-coding RNA, such as microRNAs (miRNA), small nuclear and nucleolar RNAs (snRNAs/snoRNAs), and transfer RNAs (tRNA) (Djebali et al., 2012). Present annotations now include many tens of thousands of this expanding ragtag group of RNAs, in some cases substantially outnumbering protein-encoding messenger RNAs (mRNA) (Iyer et al., 2015).

The vast majority of lncRNAs are transcribed by RNA polymerase II (pol II), as well as 5’-capped, spliced and polyadenylated (Guttman et al., 2009). The defining characteristic of lncRNA, the absence of a translated open reading frame, has received intense scrutiny given their extensive polyribosomal association and detection in ribosome profiling experiments (Banfai et al., 2012; Calviello et al., 2015; Guttman et al., 2013; Ingolia et al., 2014; van Heesch et al., 2014). Many exhibit tissue-specific expression, leading to speculation that they are subject to regulation (Cabili et al., 2011; Derrien et al., 2012). However, pol II transcription is a low-fidelity process (Struhl, 2007), which constrained by cell-type specific chromatin architecture and transcription factor expression can generate condition-specific “transcriptional noise” that might be interpreted as spatiotemporally restricted expression patterns. Furthermore, non-coding transcripts originating from regulatory regions have shown to be indicative of their activation status. Many such transcripts have been annotated as lncRNA, but they are largely actively degraded by nuclear surveillance mechanisms and as such unlikely to have any functions as RNA in *trans* (Andersson et al., 2014).

Only a small proportion of human lncRNA have known regulatory functions (Quek et al., 2014). It remains unclear which lncRNA are likely to be functional as distinct RNA species, either by encoding a short peptide or directly through interactions with proteins and nucleic acids (Rinn and Chang, 2012); which ones may be non-functional byproducts of a stochastic cellular environment; and which ones may indicate or influence transcriptional activity via the process of transcription itself but not as functional RNA (Ulitsky and Bartel, 2013). Attempts to classify lncRNA rely on either conservation, chromatin signals, or most often the genomic position and orientation relative to coding genes (extensively reviewed by Laurent et al. (2015)). However, the heterogeneity in the form and function of lncRNAs remains a major obstacle making it difficult 1) to prioritize lncRNAs for in depth characterization and 2) to generalize knowledge derived from individual lncRNAs to other lncRNAs.

RNAs that share common steps of RNA biogenesis and maturation often have similar functions as in the case of miRNAs, snRNAs and tRNAs. The idea is generalizable to mRNAs as well assuming common RNA metabolism behavior reflects common regulation by trans-acting factors, which have been shown to coordinate the expression of mRNAs encoding functionally related proteins (Keene, 2007). Since lncRNAs (and small non-coding RNAs) do not encode a separate molecule and are themselves the final active cellular molecule, their biogenesis and maturation necessarily constrain the types of regulation the participate in. We propose lncRNAs exhibiting similar RNA metabolism behaviors may have similar functional capacities. Therefore, we systematically profiled RNA metabolism at multiple steps across all RNAs in the human genome. We quantitatively examined each major step, specifically transcription, splicing, degradation, localization and translation, and found substantial differences between annotated lncRNA and mRNA. For instance, computational modeling of individual intron excision revealed mechanistic differences in splicing between lncRNA and mRNA introns. Our observations motivated us to perform a new annotation-agnostic, unsupervised classification of RNAs based on their full RNA metabolism profile. Each of seven classes contain both mRNAs and lncRNAs to a different extent, exhibit distinct evolutionary patterns and fitness constraints, differential sensitivity to cellular RNA regulatory pathways and different relationships between steps of RNA metabolism, significantly expanding the current definition based on coding potential only: Both lncRNAs and mRNAs exist on a spectrum of biogenesis and maturation, and can be grouped into behaviorally-coherent subclasses that will be invaluable for detailed functional inference.

## Results

### Differences in expression and maturation between coding and non-coding genes

Unlike mRNAs, for which a single copy can be translated numerous times, a lncRNA must be expressed at sufficient levels for the RNA product to have a function in a given cellular context. Therefore, we performed strand-specific paired-end RNA sequencing in triplicate using ERCC spike-in RNAs in human embryonic kidney cells (average depth of ∼31 million uniquely aligned reads) to determine average RNA copy numbers per cell for 27,803 genes (Supplementary Fig. 1a-c). Typical of RNA-seq experiments, we detected a low expression regime (n=13,685 genes) and a high expression regime (n=14,118 genes) with average RNA copies per cell of 0.05 and 11.45, respectively. Protein-coding genes exhibited higher expression than lncRNAs and pseudogenes (Fig. 1a), a general observation confirmed by examination of 101 different tissues and samples from ENCODE (Supplementary Fig. 1d). Only 20% of lncRNAs were in the abundant population compared to 68% of coding genes. Since the presence of an intron can enhance expression (Hir et al., 2003), it was not surprising that only 2.6% of protein coding genes in the high expression regime were intronless, compared to 32% and 51.9% of lncRNAs and pseudogenes, respectively (Supplementary Fig. 1e, f). Yet, multi-exonic lncRNAs exhibited were less robustly processed (Fig. 1b, see Supplement for estimation of primary and mature RNA expression). These results clearly demonstrated and confirmed differences in steady-state mature RNA levels and RNA maturation between coding and non-coding RNA (Cabili et al., 2011). Furthermore, most lncRNA were not expressed at a copy number to be consistent with a *trans* -acting function for the RNA product. Since RNA copy number estimates were derived from a population of cells, we cannot exclude the possibility that low copy RNAs represent high expression in a minor subset of cells. However single-molecule imaging of numerous lncRNAs in HeLa cells found that low expression was not explained by such ‘jackpot’ cells (Cabili et al., 2015).

**Figure 1.**
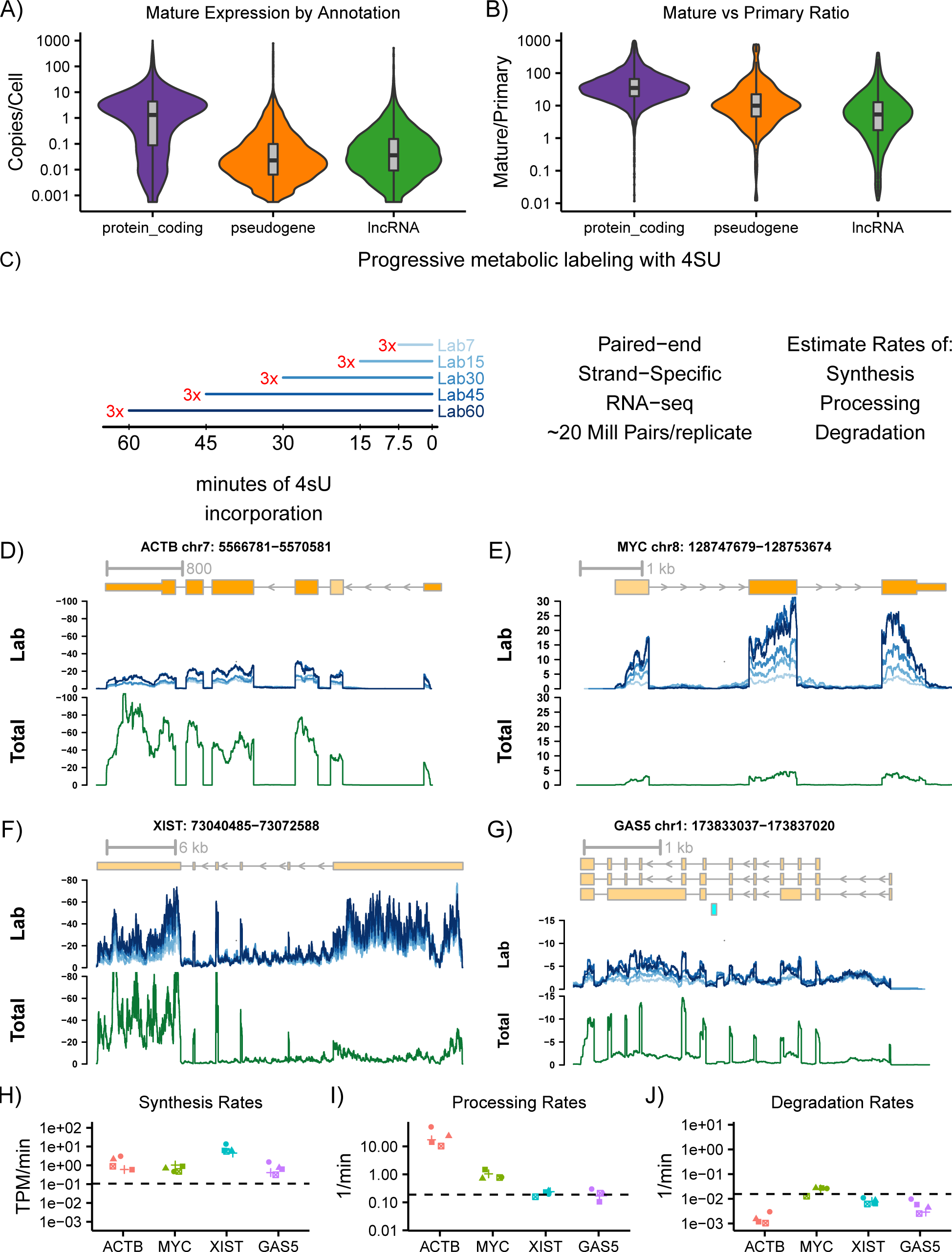
Progressive metabolic labeling of RNA. The average RNA copy number per cell (CPC) for endogenous genes was estimated using the fit determined for the ERCC spike-in RNAs. (a) CPC distribution of coding, pseudogene and lncRNA genes. (b) The ratio of mature CPCs versus primary CPCs for the high expression population of genes based on annotation category. (c) Time-points of progressive 4SU labeling. Longer labeling correspond to darker shades of blue. Coverage (library size normalized fragments per million) of 4SU data (blue) and total RNA (green) profiles for (d) *ACTB*, (e) *MYC*, (f) *XIST*, and (g) *GAS5*. Rates of (h) synthesis, (i) processing, and (j) degradation of *ACTB*, *MYC*, *XIST*, and *GAS5* for each of the five labeling times calculated using INSPEcT. The y-axis represents the full range of values for that rate and the dashed line is the average.

### Progressive metabolic labeling of RNA

To obtain a mechanistic understanding of the differences in behavior between mRNA and lncRNAs, we generated progressive snapshots of RNA production, maturation, and degradation by metabolic labeling of RNA with 4-thiouridine (4SU) (Windhager et al., 2012). After pulse-labeling cells with 4SU for 7.5, 15, 30, 45, and 60 minutes, total RNA was purified and newly-transcribed/labeled RNA was subsequently biochemically separated and strand-specifically paired-end sequenced in biological triplicate (Fig. 1c). The 4SU-labeled samples were sequenced to an average depth of 18 million uniquely mapping read pairs.The fraction of primary transcript in 4SU samples was high relative to other HEK293 data types representing different stages of RNA maturation, including genomic run-on (GRO-seq) (Fong et al., 2014) and RNA-seq of cellular fractions (nuclear, cytoplasm, cytosol, polyribosomal (Sterne-Weiler et al., 2013; Sultan et al., 2014)) (Supplementary Fig. 1g). We detected coverage for 79.6 million nucleotides in the 4SU-samples; by comparison, 171.7 million nucleotides of the human genome are annotated across all GENCODEv19 (Harrow et al., 2012) (Supplementary Fig. 1h). There was a strong correlation between 4SU and GRO-seq for genes detected in common (R = 0.83). Unstable regions, such as introns of coding genes and lncRNAs (Supplementary Fig. 1i-m), exhibited the most pronounced coverage differences between the two methods. Overall, 4SU-labeling was comparable to GRO-seq for analysis of RNA synthesis but did not require *in vitro* perturbation of nuclei.

Scrutiny of individual loci revealed increases in production of mature mRNA from 7.5 to 60 minutes for well-transcribed genes *ACTB* and *MYC* (Fig. 1d, e). Since *ACTB* had higher steady-state levels than *MYC*, we could correctly infer that *ACTB* mRNA was more stable than *MYC* mRNA. We also detected progressive increases in production of the relatively stable lncRNAs *XIST* and *GAS5*, though they exhibited less complete intron excision (Fig. 1f, g). We inferred synthesis, processing, and degradation rates for genes transcriptome-wide by comparing primary and mature RNA concentrations of 4SU-labeled RNA to total RNA using INSPEcT, a differential equation-based modeling framework (de Pretis et al., 2015). The inferred rates of synthesis, processing, and degradation from different labeling times were precise and consistent with the observed behavior (Fig. 1h-j). The two lncRNAs exhibited slower splicing of individual introns and lower gene-level processing rates than the two coding genes suggesting fundamental differences in splicing mechanisms.

### Splicing differences between coding and non-coding genes

Together with previous studies indicating lncRNAs are less efficiently spliced (Tilgner et al., 2012), the differences we observed for individual intron removal for specific lncRNAs prompted us to analyze splicing differences between individual lncRNAs and coding introns in detail. For each junction with sufficient coverage across all labeling time-points (n = 21,782), we calculated an intron-centric splicing value, θ (Supplementary Fig. 2a), which ranged from 0 (unspliced) to 1 (spliced). We identified three clusters of introns representing fast, medium, and slow intron excision dynamics (Fig. 2a). Introns of lncRNA were 17.3x more likely than introns of coding genes to belong to the slow class (Fig. 2b). Additionally, the slow class was 8.3x more likely to contain exons exhibiting higher skipping (low spliced-in values, Ψ, see supplemental methods) (Fig. 2c). Mirtrons were spliced out more quickly than introns not containing small RNAs, while snoRNA-containing introns were spliced more slowly (Fig. 2d).

**Figure 2.**
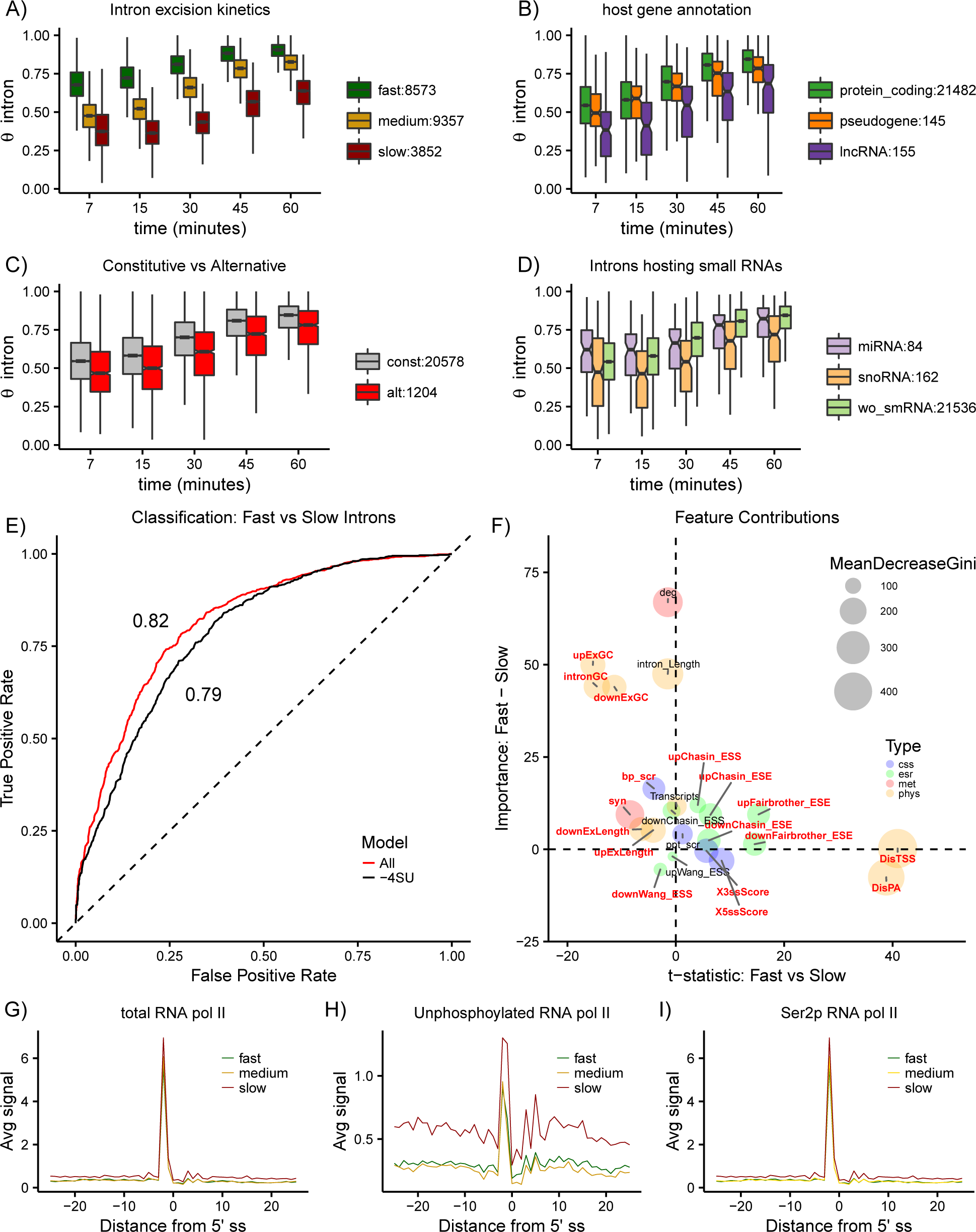
Dynamics of intron excision. Introns with fast, medium and slow intron excision speeds were were identified by clustering θ values of introns with sufficient data (n=21,782) at all labeling time-points.Violin plots depicting the distribution θ of values for introns grouped based on (a) excision dynamics, (b) host gene annotation, (c) constitutive or alternatively spliced adjacent exons (mean Ψ values =0.37), and (d) the type of small RNA hosted by the intron. (e) The average receiver operating characteristic (ROC) curve for predictions using 5-fold cross-validation using all features (black) or all features excluding those derived from 4SU data such as synthesis and degradation rates (red). (f) Bubble plot depicting the differential contribution and importance of features to classification. The difference (fast-slow) in mean decrease in model accuracy for each class (y-axis) plotted against the t-statistic of the difference of means between intron classes (fast-slow). The circle size is the mean decrease in Gini coefficient representing the importance of that feature for the classification. Features with statistically significant differences (t-test, p < 0.05) are labeled in bold red text. The average NET-seq signal +/ -25 nucleotides from the 5’ splice site of (g) total RNA polymerase II, (h) unphosphorylated RNA polymerase II, and (i) ser2p RNA polymerase II for fast (green), medium (yellow), and slow (red) introns.

To investigate the mechanisms underlying the different splicing behavior, we used features corresponding to nucleotide composition and length, canonical splicing signals, and exonic splicing regulatory elements as well as RNA metabolism (synthesis and decay on the gene-level), in a random forest classifier trained to discriminate between the fast and slow intron classes (Supplementary Fig. 2b). The model was trained using all features as well as separately excluding rates derived from the 4SU data (synthesis and degradation rates). Both models had similar performance, with a modest improvement when adding metabolic features (auROC from 0.79 to 0.82; Fig. 2e). Similar features were important using an orthogonal approach using individual splicing models to predict for coding introns, lncRNA introns, snoRNA host introns, and mirtrons using random forest regression (Supplementary Fig. 2c-f).

The distance of introns from the transcription start site (TSS) and, especially, to the transcription end site (TES) were important physical features for prediction and positively correlated with splicing speed (Fig. 2f). The GC-content of the introns and flanking exons were of particular note because they were more important for the prediction of fast introns and significantly lower for fast introns and flanking exons. With respect to canonical splicing signals, fast introns exhibited stronger splice sites, weaker branchpoints but similar polypyrimidine tract scores. Upstream and downstream exons flanking fast introns exhibited significantly higher levels of ESEs and lower levels of ESSs compared to slow introns, consistent with recent evidence for purifying selection to preserve exonic splicing signals in lncRNAs (Haerty and Ponting, 2015; Schuler et al., 2014).

The higher synthesis rates of genes containing slow introns (Fig. 2f, “syn”) prompted us to examine phosphorylation status of the C-terminal domain (CTD) of RNA polymerase II (Hsin and Manley, 2012) at splice sites using native elongating transcript sequencing data (NET-seq) (Nojima et al., 2015). We observed similar total and ser2p NET-seq signal at the 5’ splice site (Fig. 2g,i), but substantially more unphosphorylated RNA poll II signal at flanking 5’ splice sites of slow introns (Fig. 2h). These data indicated the relative absence of proximal RNA polymerase II phosphorylation as a potential mechanism for the decreased splicing efficiency of these lncRNA introns, consistent with *in vitro* results (Hirose et al., 1999). In combination, GC-content, splicing regulatory elements and RNA pol II phosphorylation were very different between fast and slow introns and were important for their classification, as well as for discriminating coding and lncRNA introns.

### Differences in RNA metabolism between coding and non-coding genes

The differences in steady-state transcript levels observed between lncRNA and mRNA must be due to differences in the metabolism of lncRNAs. To monitor the full sequence of RNA life, from synthesis to translation, we complemented the 4SU-derived features with the following quantitative estimates of subcellular localization and translation status all in HEK293 cells (see Supplement for detailed explanation): the enrichment of cytosolic over nuclear expression (CytN) (Sultan et al., 2014); the enrichment of polyribosomal over cytosolic expression (PlyC) (Sterne-Weiler et al., 2013); and translational potential (TrP), which represents the amount of translating ribosomes on a transcript based on their codon-by-codon movement (Calviello et al., 2015). We focused the rest of our analysis on the 15,120 genes for which we had complete data.

We observed high correlation between rates calculated from different labeling times (Supplementary Fig. 3 a-d), and degradation rates were similar to previous estimates (R=0.53) (Gregersen et al., 2014). Consistent with earlier results from mouse cells (Rabani et al., 2011), transcripts encoding ribosomal proteins exhibited high synthesis and processing as well as low degradation rates, confirming the quantitative behavior of the estimated rates (Fig. 3a). Very different RNA metabolism patterns generated similar steady-state behavior: for example, KRAB domain transcription factors exhibited average synthesis rates and high degradation rates, while extracellular proteins exhibited low synthesis rates and average degradation rates (Fig. 3a). Yet, these two groups showed similarly low steady-state RNA levels, revealing the limitations of solely assaying abundance.

**Figure 3.**
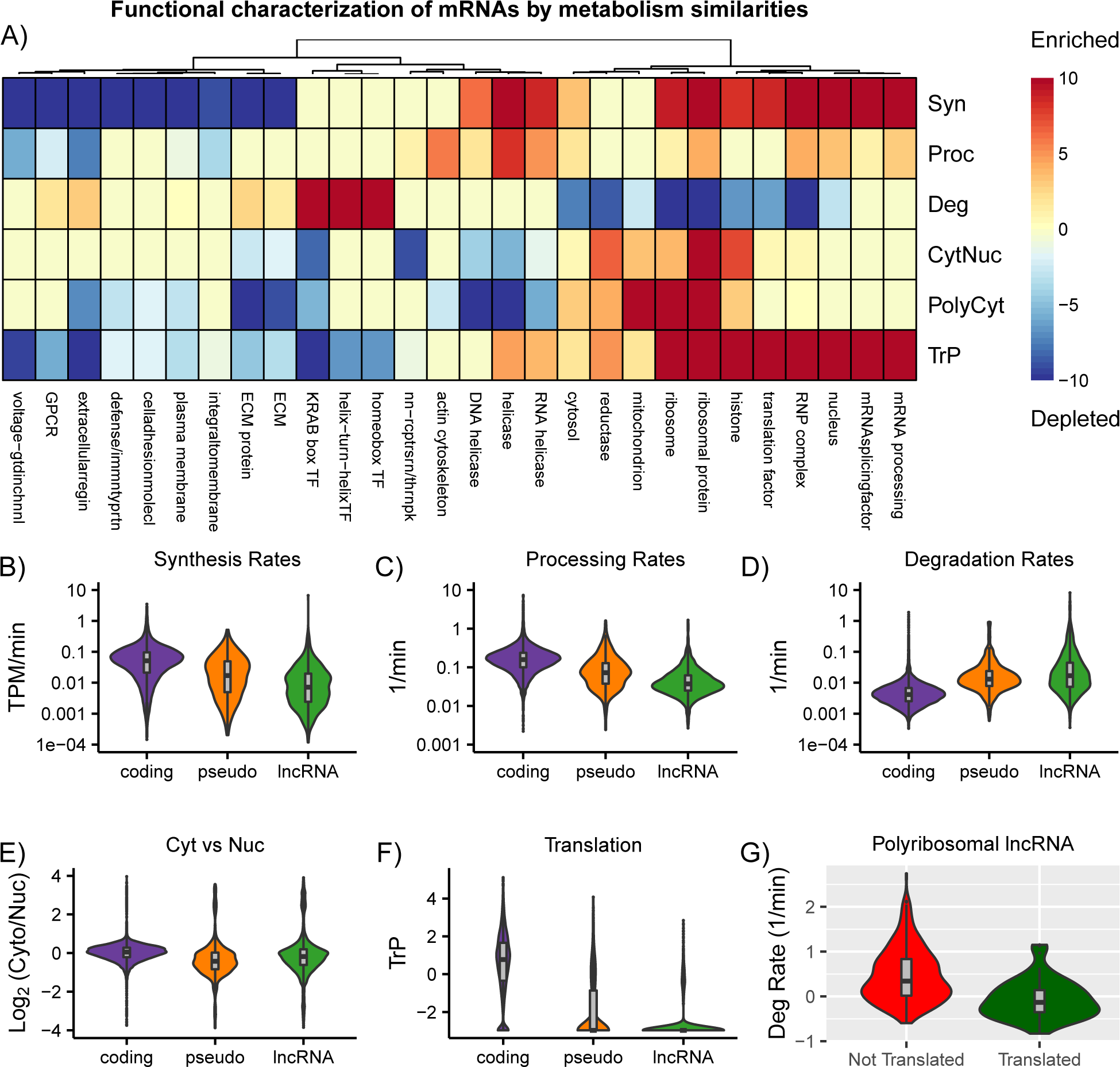
RNA metabolism of mRNA and lncRNA. (a) Clustering of gene ontology enrichment for protein-coding genes based six hallmarks of RNA processing: synthesis rates (Syn), processing rates (Proc), degradation rates (Deg), cytoplasmic vs nuclear localization (CytNuc), polyribosomal vs cytoplasmic localization (PolyCyt), and translational status from ribo-seq data (TrP). Each feature (e.g. Syn) was rank-ordered and the p-value of enrichment for each gene sets was calculated using PantherDB. The scale indicates the log_10_(p-value) for enriched gene sets (red) and -log_10_(p-value) for depleted gene sets (blue). (b-g) Distribution of (b) synthesis rates, (c) processing rates, (d) degradation rates, (e) cytoplasmic vs nuclear localization, (f) translation for coding, lncRNA, and pseudogenes. (g) Distribution of degradation rates for polyribosomal lncRNA (PolyCyt > 0.1) divided into groups based on the presence or abscence of a RiboTaper-detected translated ORF from ribosome profiling data in HEK293 cells for coding, lncRNA, and pseudogenes.

The average synthesis and processing rates for lncRNAs were 3.4x and 2.9x lower than protein coding genes, respectively (Fig. 3b, c). LncRNAs had 9.6x higher average degradation rates than mRNAs (Fig. 3d), a conspicuous difference to earlier less comprehensive studies reporting lncRNAs to be on average only 1.6x (Clark et al., 2012) or 0.97x (Tani et al., 2012) less stable than mRNAs. Overall lncRNAs were only slightly more localized in the nucleus than mRNAs (Fig. 3e) and their polyribosomal localization was similar (Supplementary Fig. 3e).The vast majority of lncRNA did not contain an actively translated open reading frame (ORF) (Calviello et al., 2015)(Fig. 3f), confirming their status as *bona fide* non-coding RNA. Amongst polyribosomal lncRNA, those exhibiting evidence for translation were more transcribed (Supplementary Fig. 3f) and more stable (Fig. 3g).

### Annotation-agnostic classification of genes using RNA metabolism profiles

Although we observed substantial differences in numerous aspects of RNA metabolism between annotated coding genes and lncRNAs, we also noticed that both lncRNA and mRNA exhibited a wide and overlapping range of behavior (Fig. 2,3). This highlighted the heterogeneity in metabolism of both lncRNA and mRNA with respect to annotation. We hypothesized that genes with similar RNA metabolism features would reflect the coordinated activity of specific RNA processing factors. Importantly, for non-coding RNAs this classification would help distinguish which, if any, of the wide range of regulatory mechanisms they might participate in. Therefore, we clustered all 15,120 genes (including annotated coding and non-coding as well as pseudogenes) *de novo* based on all six features of the metabolism profiles. The number of clusters was determined using the gap-statistic (Supplementary Fig. 4a).

We identified seven RNA classes containing from 921 to 3293 genes (Fig. 4a). Classes c1-c4 were enriched for coding genes (n=10,793), although they contained 220 lncRNAs (Fig. 4b). Classes c5-c7 were enriched for lncRNA (n=1752), although they contained a similar number of coding genes (n=1838). Importantly, all GENCODE-defined lncRNA subcategories (e.g., lincRNA or antisense) were non-specifically found across clusters c5-7 (Fig. 4c), demonstrating that classification based on positional genomic features does not coincide with specific lncRNA behavior. Consistent with earlier studies that indicated many processed-transcript RNAs to be translated (Calviello et al., 2015), this biotype was more likely to be in c1-c4 than other lncRNA biotypes.

**Figure 4.**
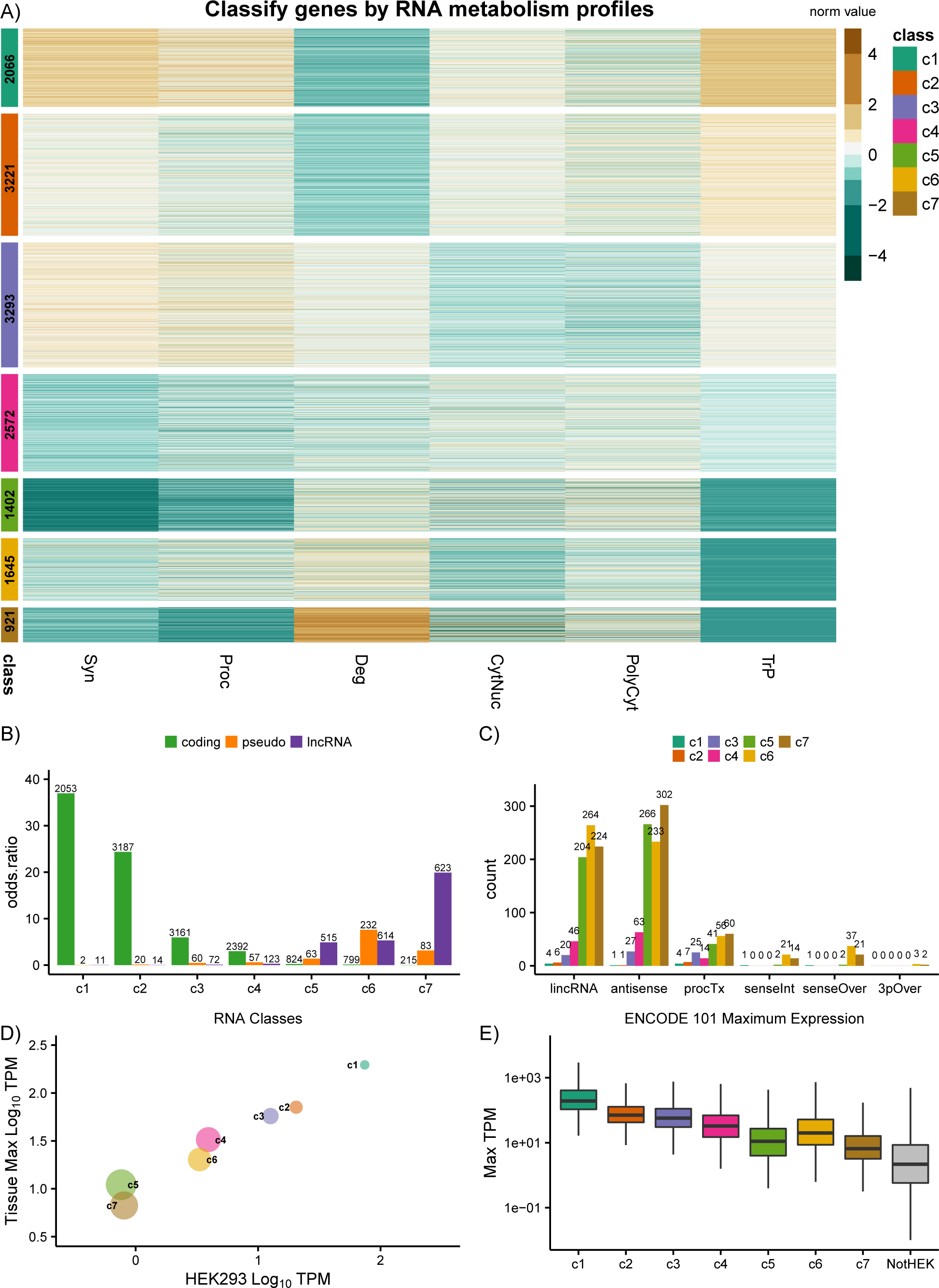
Classification of genes by RNA metabolism profiles. (a) Centered and scaled values for each metabolic feature for each of seven classes determined by k-means clustering. The number on the class labels represents the number of genes in each class. (b) Odds-ratio (Fishers exact test) for overlap between class membership and GENCODE V19 annotation classes. Numbers of genes in each class and annotation category shown above bars. (c) The number of lncRNA genes a given GENCODE V19 biotype for each RNA class. Numbers of genes in each class and biotype category shown above bars. (d) Bubble plot of median value of HEK293 expression (x-axis) and of maximum TPM across 101 tissues/cell lines from ENCODE (y-axis) for all genes in a class. The bubble size represents tissue specificity score (higher score = more tissue restricted). (e) Boxplot showing distribution the maximum TPM across 101 ENCODE cell lines and tissues for genes in each RNA class and for genes excluded from the clustering of RNA features due to insufficient data in HEK293 cells.

Even though protein-coding genes were enriched in classes c1-c4, they occurred in all seven classes enriched for distinct characteristics and encoding functionally related proteins (Supplementary Fig. 4b). Genes in c1 and c2 exhibited the most similar RNA metabolism profiles, with c1 genes having modestly higher synthesis and processing rates, lower degradation rates and higher translation. The c1 genes exhibited the highest expression and lowest tissue specificity, consistent with “housekeeping”/constitutive genes such as mRNAs encoding ribosomal proteins. Genes in c3 were synthesized and processed well, but had relatively higher degradation rates and were less cytoplasmic; this class was enriched for transcription factors, particularly the aforementioned KRAB domain transcription factors. These genes may be subject to nuclear retention as a mechanism to reduce cytoplasmic gene expression noise created by transcriptional bursts (Battich et al., 2015; Halpern et al., 2015); indeed, c3 genes exhibited a lower level of cytoplasmic localization relative to other classes in a recent study of subcellular expression in mouse liver (Supplementary Fig. 4d). Genes in c4 had relatively lower synthesis, processing and translation rates along with higher degradation rates compared to c1-3 and were enriched for receptors; they were also depleted of ribosomal proteins and RNA processing factors. These genes exhibited the most tissue specificity of c1-4 and had steady-state expression levels similar to genes in c6. Within c5-7, the c6 genes had the highest synthesis rates and most nuclear localization and were enriched for pseudogenes along with calcium and ion channel-encoding mRNAs. Genes in c5, typified by GPCRs and ion channels, and c7, typified by peptide hormones, exhibited similar steady-state levels of expression but very different degradation rates and localization. The c7 genes had the highest proportion of protein-coding genes with no functional classification and that are part of the “missing” proteome (Wilhelm et al., 2014) (Supplementary Fig. 4c), raising the possibility that many of these genes may not (or no longer) be protein-coding.

The classes exhibited overlapping gene expression distributions (Supplementary Fig. 4e), and thus were not recapitulated by steady-state expression or any individual RNA metabolism feature. For all classes, the maximum expression level of genes across tissues/cell-lines (101 tissue/cell-lines from ENCODE) were correlated with their expression in HEK293 cells (Fig. 4d, see Supplementary Methods for details). Classes c4-c7 exhibited the most tissue/cell-line specific expression (Supplementary Fig. 4f). Additionally, genes not expressed in HEK293 cells had the lowest distribution of maximum expression levels across tissues/cell-lines (Fig. 4e). These results supported the generality of RNA metabolism-derived classes and not simply a HEK293 cell-specific behavior. Importantly, they highlighted the distinct utility of our new classes compared to existing classifications or biotypes (Supplementary Fig. 4g).

### Classes exhibit specific evolution and fitness signatures

Our RNA metabolism-derived classes exhibited differences in gene age (origination in vertebrate phylogeny derived from (Zhang et al., 2010)). Classes c1, c2, and c3 were enriched for ancestral protein-coding genes predating the divergence of vertebrates (Fig. 6a). Most lncRNAs originated throughout mammalian and primate evolution, while pseudogenes were mainly gained in primates. Genes in c4 and c5 were gained prior to the divergence of eutherians mammals. Classes c3, c6 and c7 showed significant enrichment for gains in the primate lineage, with the latter two enriched for human-specific genes. These two classes had the highest degradation rates, which more strongly correlated with gene age than synthesis rates (Fig. 5b, c). This is consistent with observations of high turnover of lineage-specific lncRNAs in tetrapods (Necsulea et al., 2014) as well as rodents (Kutter et al., 2012), specifically. Unique to our approach, we were able to observe that young genes have managed to be synthesized but not avoided being degraded, as has been suggested for the balance between splicing and polyadenylation of RNAs produced from divergent transcription (Wu and Sharp, 2013).

**Figure 5.**
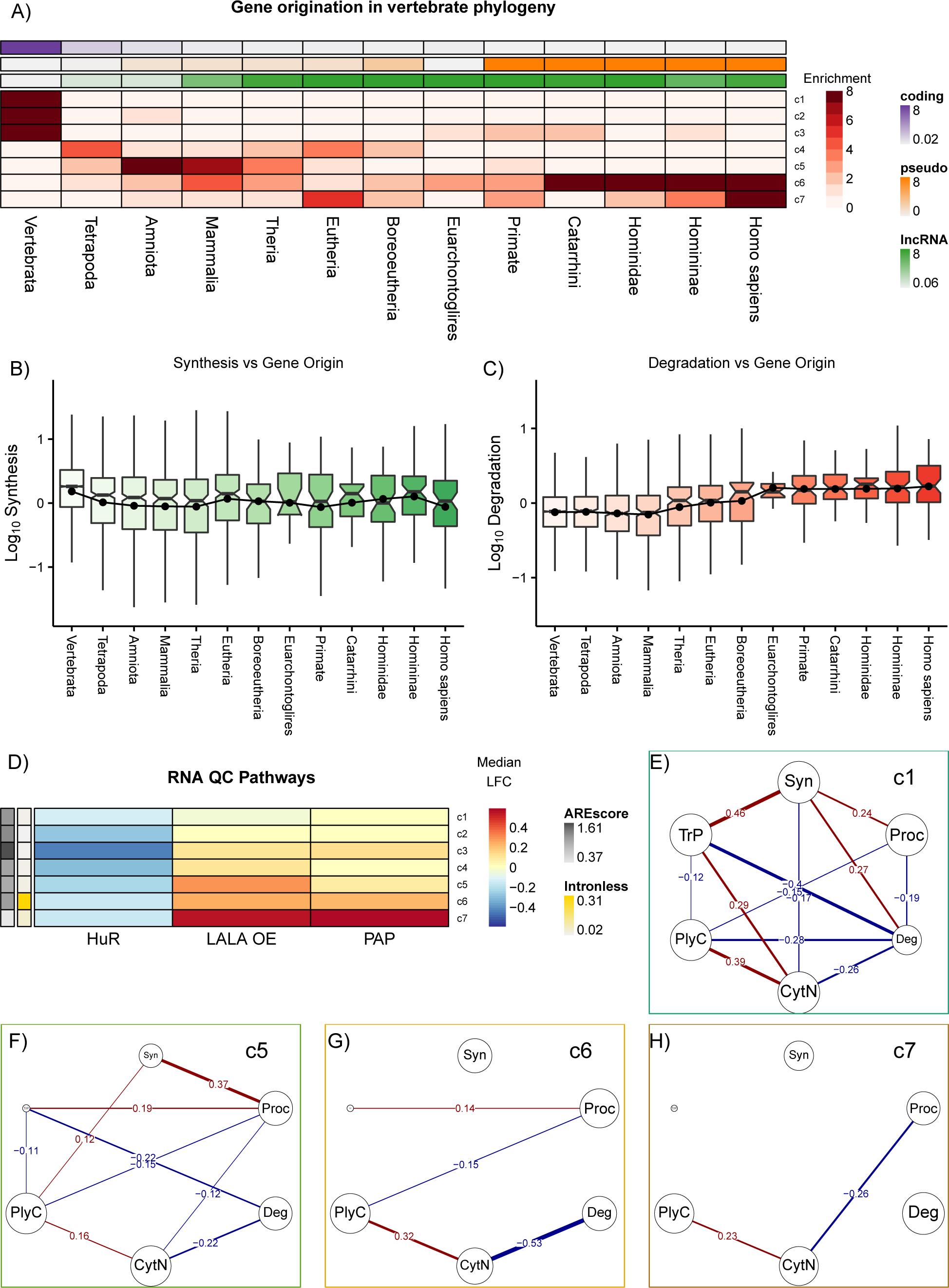
Evolutionary and regulatory differences between RNA classes. (a) Heatmap of -log10 p-value of overlap enrichment between the phylogenetic point of gene origination and RNA classes. Odds-ratio for overlap between gene origination and annotation category, specifically coding (purple), pseudogene (orange), or lncRNA (green). Boxplot of (b) synthesis and (c) degradation rates for genes grouped by origination class. A line is depicted connecting the means for each class (point). (d) Log 2 fold change of RBP perturbation - control in HEK293 cells. Median AREscore of all 3’UTR sequences from class (grey) and percent intronless (yellow). (e-h) Partial correlation coefficient between different RNA metabolism features c1, c5, c6, and c7.

### Distinct regulatory pathways shape RNA classes

The classes responded in specific ways to perturbations of cellular RNA quality control and/or regulatory pathways (Fig. 5d). Genes in c3 were the most down-regulated upon depletion of HuR/ELAVL1, an RBP antagonizing AU-rich element (ARE) mediated decay. The genes’ nuclear preference is thus aligned with the pre-mRNA stabilizing function of HuR (Mukherjee et al., 2011). They also exhibited the highest 3’ UTR ARE content and degradation rates except for c7, consistent with a particular sensitivity of these genes to cytoplasmic ARE-mediated decay mechanisms. The nuclear poly A binding protein (*PABPN1*) and polyA polymerase (*PAP*) mediated RNA decay (PPD) pathway limits the accumulation of inefficiently processed nuclear RNAs (Bresson et al., 2015). Genes in c5 and especially c7 exhibited the slowest processing rates and increased the most upon over-expression of a dominant-negative *PABPN1* mutant (LALA) that binds RNA but cannot stimulate *PAP* (Bresson et al., 2015). Interestingly, only c5 and c7 genes exhibited strong negative correlation between processing rate and export to the cytoplasm (Fig. 5 f,h), explaining the lower sensitivity of c6 genes to PPD. Furthermore, the classes exhibited unique patterns of response to depletion of different RBPs in K562 cells from ENCODE (Supplementary Fig. 5a). Altogether these results indicated that RNA classes are controled by distinct regulatory pathways in multiple human cell types.

We examined the relationship between different steps of RNA metabolism by quantifiing each pairwise association between any two features while accounting for all other features (partial correlation analysis, see Supplementary Methods). For example, class c1 genes exhibited the most interdependence between different steps of RNA metabolism (Fig. 5c), while c6 and c7 (Fig. 5 g,h) exhibited the least interdependence. The coupling between various steps of RNA metabolism is expected to be dependent on cis-regulatory elements allowing these genes to interact with the responsible RNA processing machinery. Thus, less coupling of RNA metabolism is indicative of classes enriched for “younger” genes (Fig. 5a), with a lower fraction of sequence under evolutionary constraint.

### lncRNA in different classes exhibit different behavior

Finally, we focused our characterization of classes on lncRNAs, specifically asking which classes were enriched for known functional lncRNAs from lncRNADB v2.0 (Quek et al., 2014). Classes c1, c2, and c3 exhibited the strongest overlap with lncRNADB (Fig. 6a) compared to c4 and c6, which were only ∼1.5x more likely to be found in lncRNADB than expected. In contrast, c5 and especially c7 were less likely to be in lncRNADB than expected. Importantly, lncRNA biotypes showed fewer distinct differences in overlap with lncRNADB, with processed transcripts, as expected, showing the strongest enrichment (Supplementary Fig. 6a).

**Figure 6.**
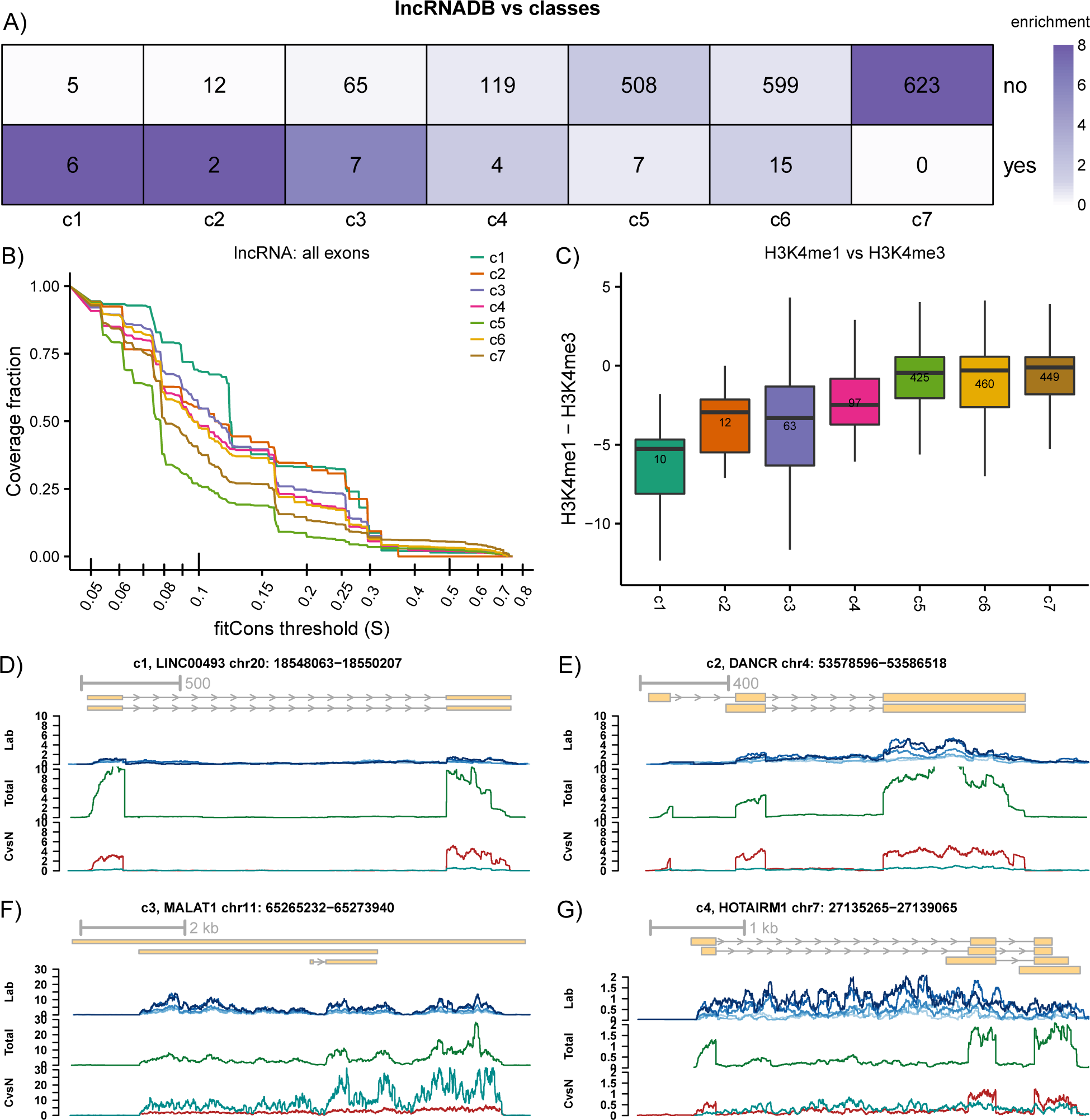
Distinct behavior of lncRNA classes. (a) The odds-ratio of the overlap between lncRNAs that were either found (“yes”) or not found (“no”) in lncRNADb for each RNA class. The numbers represent the gene count in each category. (b) The fraction of nucleotides of lncRNA exons in a particular class with a fitCons score > S calculated. (c) Boxplot of the distribution of H3K4me1 vs H3K4me3 input normalized ChIP-seq signals in the promoter (-250 to +750 nucleotides from the annotated transcriptional start site) of genes in each class. (d-g) Coverage of 4SU (blue), total (green), cytoplasmic (red) and nuclear (cyan) RNA profiles for lncRNAs belonging to c1-c4.

Although informative the gene origination analysis suffered a lack of comprehensiveness particularly for lncRNA genes. Thus, we compared genic regions using FitCons, an approach that integrates sequence divergence in primates and sequence polymorphisms across human poulations together with functional genomic data to estimate the probability that a point mutations will have a fitness consequence (Gulko et al., 2015). Independent of the gene region (cds exon, coding exon) or annotation (coding/lncRNA) examined, c1-4 and c6 had higher fitCons scores than c5 and c7 (Fig. 5b and Supplemental Figure 5c, d). Of the classes enriched for lncRNAs (c5-7), c6 had better coverage scores and behaved more like the the “coding-like” c1-4 genes. c5 lncRNA exons had lower fitCons coverage than those from c7, which have substantially higher degradation rates. The lower constraint on c7 (younger) exonic lncRNA sequence was consistent with earlier evolutionary studies (Kutter et al., 2012; Necsulea et al., 2014). These differences for “fitness consequences” in the exonic sequence encoded by both coding genes and lncRNAs strongly support the utility of our classes, in general, and with respect to lncRNA biotypes, in particular (Supplementary Fig. 5e).

Histone modifications at transcription start sites (TSSs) have been used to distinguish between promoter (H3K4me1) and enhancer-like regulatory elements (H3K4me3). This approach has been used to define different classes of intergenic lncRNAs (Marques et al., 2013). We compared the ratio of H3K4me1 versus H3K4me3 at annotated TSSs (-250 to +750 bp) of lncRNAs in different classes (Fig. 6c). We found that c1-c4 were had the lowest H3K4me1:H3K4me3 ratio and c5-c7 had the highest ratio. The c1-c4 lncRNAs are thus not likely to have potential for enhancer-like cis regulation of neighboring genes, compared to c5-c7. The latter classes, particularly c7, may involve a mechanism indicative of transcriptional regulatory activity but unlikely to function as RNA.

The classification of lncRNAs based on their RNA metabolic pattern thus provided insights into their potential functions. For example, an unstable nuclear lncRNA cannot perform a cytoplasmic trans-regulatory function. *LINC00493* (Fig. 6d) and *DANCR* (Fig. 6e) were similar with regards to synthesis, processing, degradation and export, but belonging to different clusters. However, *LINC00493*, belonging to c1, contained a highly translated ORF with peptide evidence, thus providing additional evidence to annul its status as a *bona fide* lncRNA (Michalik et al., 2014). *DANCR* was a member of c2 and has been reported to be involved in the antagonism of miRNAs (Kretz et al., 2012; Yuan et al., 2015), consistent with its high stability and cytoplasmic localization. *MALAT1*, a well-known nuclear lncRNA, belonged to the category of stable and nuclear localized c3 genes (Fig. 6f), consistent with its known role in splicing (Tripathi et al., 2010). The genes in c4 and c6 had similar synthesis, processing, and degradation rates, but c6 genes were strongly localized to the nucleus. Accordingly, *TUG1*, a lncRNA capable of recruiting polycomb-repressive complex 2 and silencing a subset of genes (Khalil et al., 2009), was a member of c6 (Supplementary Fig. 6c). The lncRNAs in c4 may have cytoplasmic and/or nuclear functions. *HOTAIRM1A*, one of these genes, has a nuclear function given its ability to modulate the expression of neighboring HOX genes (Zhang et al., 2009), although we also detected a spliced cytoplasmic isoform (Fig. 6g). The lncRNAs in c5 and especially c7 were most likely quickly degraded transcriptional byproducts, and thus are the least likely to be functional as RNAs (Supplementary Fig. 6b, d). In this way, representative well-studied lncRNA now provide specific hypotheses for additional lncRNA belonging to the same cluster.

## Discussion

This study represents one of the most multi-faceted and quantitative examination of RNA metabolism in a human cell line. Consistent with other studies in human we also observed that lncRNAs were slightly more enriched in the nucleus compared to mRNA (Cabili et al., 2015; Derrien et al., 2012; Djebali et al., 2012). However their remains a popular misconception regarding strong preference toward nuclear localization for lncRNAs (Ulitsky and Bartel, 2013), which may be due to chromatin-associated regulatory activity of a few of the most well known lncRNAs, such as *XIST* or *MALAT1*. The relatively poorer processing and higher degradation of lncRNAs would at least partially explain the less cytoplasmic/more nuclear localization of lncRNAs relative to mRNAs rather than specific regulation of lncRNA localization. Additionally, we confirmed that steady-state lncRNA copy numbers in human are typically ∼10x lower than mRNA (Cabili et al., 2011) and show this is due to differences in both synthesis and degradation. This is an important question that cannot be answered from steady-state RNA levels alone. One could envision even a low abundance lncRNA having a *trans*-acting function if it is stable and it may even participate in sequential interactions thereby “amplifying” its regulatory capacity similar to mRNAs being translated multiple times.

We are aware of only one study that explicitly compared mRNA and lncRNA stability in human cells, which observed minimal differences in RNA stability (Tani et al., 2012); a study conducted in mouse neuroblastoma cells also observed similar stability (Clark et al., 2012). These studies used different methods for measuring expression and determining stability and may not have had sufficient depth to capture rare RNAs. Only 30 lncRNAs with measured stability are found in both datasets. These 30 lncRNAs belonged to class c1/2/3/6 and, importantly, none were in c7, the extremely unstable and most tissue-specific class accounting for 31.6% of our detected lncRNAs. Consistent with our findings, Rabani et al. (2014) also point to substantial differences between ‘lincRNA’ and mRNA stability in LPS response of mouse dendritic cells, but they reported similar synthesis and processing rates between mRNA and a subset of abundant ‘lincRNA’. Each study used different annotation information and none of these studies focused on the same group of lncRNAs or even “lincRNAs”. Beyond technical, species- and context-specific differences, the heterogeneity of lncRNAs and the lack of a coherent classification make it inherently difficult to reconcile conclusions from different studies.Thus our annotation-agnostic unsupervised classification of genes based on six hallmarks of RNA processing, represents an important step toward providing functional and relational coherence to the field.

Intriguingly, we not only identified lncRNA in protein-coding dominated classes, but also identified hundreds of protein-coding transcripts with metabolism profiles that appear more lncRNA-like. In fact, any given gene exists on a continuum within a specific cellular and evolutionary context, and can be in the process of gaining or losing specific characteristics. For instance, some of the protein-coding genes that cluster with pseudogenes or unstable lncRNAs are likely to never generate proteins, but this apparent pseudogenization may have occurred too recent to allow for accumulating sequence changes typically used to identify bona fide pseudogenes. Likewise, a small number of currently annotated lncRNAs do not just encode translated open reading frames with detectable peptides being produced, but their whole processing resembles protein-coding genes and to such an extent that they can now safely be re-annotated. However we find that most of the known functional lncRNAs are enriched in RNA classes showing evidence for translation c1-c4, though no detectable peptide. It is unclear the extent to which the translational signature relates to the functional lncRNAs and if they tend to be bi-functional genes, such as *SRA1* (Cooper et al., 2011).

Our results substantially expand the understanding of the features governing intron excision. We show that introns that took longer to be excised were more likely to be from lncRNAs and alternative exon usage events. This complements and confirms research indicating that these two groups are less likely to be co-transcriptionally spliced and/or less efficiently by comparing subcellular fractions (Pandya-Jones et al., 2013; Tilgner et al., 2012). Clearly, the slowly excised introns we detected would include some detained and/or retained introns (Boutz et al., 2015; Braunschweig et al., 2014). We and Braunschweig et al. (2014) both identify high GC-content, intron position, weak splice-site strength and increased polymerase occupancy, and we additionally show the importance of splicing enhancer density (higher for fast group) and silencer density (higher for slow group). Integration of NET-seq data show that slow introns tend to have more unphosphorylated RNA pol II which cumulatively with the above differences would reduce co-transcriptional recruitment of splicing factors. These results will aid future experimental dissection of the order and impact of these features on lncRNA splicing and function.

Our study substantiates a growing sentiment in the lncRNA field aptly pointed out by Quinn and Chang (2015): that we need to “recognize that [lncRNAs] exist on multidimensional spectra of biogenesis, form and function.” Inspired by half-a-century old research tritium labeling and sedimentation to investigate cellular RNA leading to the discovery of heterogeneous nuclear RNA (hnRNA) (Soeiro et al., 1966), we used a combination of genomic technologies and computational modeling to quantify numerous aspects of RNA metabolism and classify hnRNAs by their life history. Gene position/orientation independent attempts to classify RNAs, specifically lncRNAs, largely depend on evolutionary sequence conservation and/or comparison of chromatin modifications. Both of these approaches have yielded important insights however their utility is compromised by the overlapping and complex gene models, which our integrative approach proposed here can handle due to strand-specific nature of our assays. Furthermore, methods to address the evolutionary conservation and selection of RNA structural elements will be important for the inference of lncRNA-mediated regulatory mechanisms in spite of the computational observation that lncRNAs are folded than mRNAs (Yang and Zhang, 2014).

While this work focuses on a single human cell line we provide evidence that the observations are generalizable; there will undoubtedly be context-specific differences in behavior that will be the focus of future work. Moreover, quantifying these behavior at the transcript-level will be very important. Our behavioral classification of RNAs 1) provides a starting point for collecting genetic evidence for the function of lncRNAs (Sauvageau et al., 2013), and 2) enables potential inference of regulatory mechanisms for a specific RNA to other RNAs with similar behavior.

## Author Contributions

Conceptualization, N.M. and U.O.; Methodology, N.M. and U.O.; Software/Formal Analysis, N.M., L.C. and S.P.; Investigation, N.M. and A.H.; Visualization, N.M.; Writing – Original Draft, N.M. and U.O.; Writing – Review & Editing, L.C., S.P. and M.P.; Funding Acquisition, N.M. and U.O.; Resources, N.M. and U.O.; Supervision, N.M. and U.O.

## Acknowledgments

We declare no conflicts of interest. UO acknowledge support from an award by the US National Institutes of Health (R01-GM104962). NM acknowledge support from EU Marie Curie IIF (EU). All sequencing data have been submitted to the SRA/GEO GSE84722. Code and interactive-data visualization available a https://ohlerlab.mdc-berlin.de/software/Classification_of_human_genes_by_RNA_metabolism_profiles_130/.

